# *MFN2* Influences Amyotrophic Lateral Sclerosis Pathology

**DOI:** 10.1101/2021.10.30.466517

**Authors:** Kristi L. Russell, Jonathan M. Downie, Summer B. Gibson, Karla P. Figueroa, Cody J. Steely, Mark B. Bromberg, L. Charles Murtaugh, Lynn B. Jorde, Stefan M. Pulst

## Abstract

**Objective:** To better understand the pathology of amyotrophic lateral sclerosis, we used sequence data from patients seen at the University of Utah to identify novel disease-associated loci. We utilized both *in vitro* and *in vivo* studies to determine the biological effect of patient mutations in *MFN2*.

**Method:** Sequence data for a total of 140 patients were run through VAAST and Phevor to determine genes that were more burdened with rare, nonsynonymous variants compared to control longevity cohort. Variants identified in MFN2 were expressed in *Mfn2* knockout cells to determine if mutant MFN2 could rescue mitochondrial morphology defects. We identified additional rare, nonsynonymous variants in MFN2 in ALSdb that were expressed in knockout mouse embryonic fibroblasts (MEFs). Membrane potential was measured to quantify mitochondrial health upon mutant MFN2 expression. *mfn2* knockout zebrafish were used to examine movement compared to wildtype and protein aggregation in brain.

**Results:** *MFN2* mutations identified in ALS patients from our University of Utah cohort and ALSdb were defective in rescuing morphological defects in *Mfn2* knockout MEFs. Selected mutants showed decreased membrane potential compared to wildtype MFN2 expression. Zebrafish heterozygous and homozygous for loss of *mfn2* showed increased TDP-43 levels in their hindbrain and cerebellum.

**Conclusion:** In total, 21 rare, deleterious mutations in *MFN2* were tested in *Mfn2* knockout MEFs. Mutant MFN2 expression was not able to rescue the knockout phenotype, though at differing degrees of severity. Decreased membrane potential also argues for inhibited mitochondrial function. Increased TDP-43 levels in mutant zebrafish illustrates MFN2’s function in ALS pathology. MFN2 variants influence ALS pathology and highlight the importance of mitochondria in neurodegeneration.

## INTRODUCTION

Amyotrophic lateral sclerosis (ALS) is a rare, rapidly fatal neurodegenerative disease that is the result of the death of motor neurons in the motor cortex and spinal cord^1,2^. It is the most common motor neuron disease in adults. No effective treatments currently exist. Heritability estimates indicate that 60% of the variation in sporadic ALS (SALS) susceptibility can be ascribed to genetic factors, yet mutations in known ALS genes, such as *TARDBP* and *SOD1*, only account for ∼17% of SALS cases^3,4^. To identify novel ALS-causing genes, we analyzed rare variation and gene burden in 140 ALS cases compared to controls.

This led us to investigate variants in *MFN2*, which encodes mitofusin 2, and its potential function in ALS. Mitofusin 2 (MFN2) is a mitochondrial transmembrane GTPase with the GTPase domain in the cytosol^5^. MFN2 is necessary for mitochondrial fusion and the building of a complex mitochondrial network^6^. Tethering between two mitochondria starts with the heptad repeat (HR) domain followed by GTP hydrolysis^7^. MFN2 and MFN1 proteins on distinct mitochondria bind and form oligomers to help facilitate fusion into a single mitochondrion. The importance of MFN2 is highlighted by the fact that *Mfn2* knock-out (KO) mice do not survive past midgestation^6^. Mfn2 is also necessary for cerebellar development, as homozygous mutants display severe motor impairment^8^. Mutations in *MFN2* cause the motor and sensory neuropathy Charcot-Marie-Tooth type 2A (CMT2A, also known as hereditary motor and sensory neuropathy type II)^9^. *MFN2* mutations cause the most common form of dominant CMT type 2 (CMT2A) and are estimated to account for ∼20% of CMT2 patients (Fig1C)^10^. Symptoms include muscle weakness and atrophy, with sensory loss that usually affects peripheral structures such as the feet and hands. Symptoms usually begin in childhood. Compared to CMT type 1, CMT2A is associated with more severe, motor-predominant phenotypes with progressive loss of neuromuscular coordination at an earlier age^11^. *MFN2* heterozygous, homozygous, and compound heterozygous mutations have all been reported to cause severe early-onset axonal neuropathy, with disease sometimes occurring before the age of five years ^9,11-14^. Our findings support MFN2’s additional function in ALS pathology and highlight the importance of mitochondrial dysfunction in neurodegeneration.

**Fig 1.**
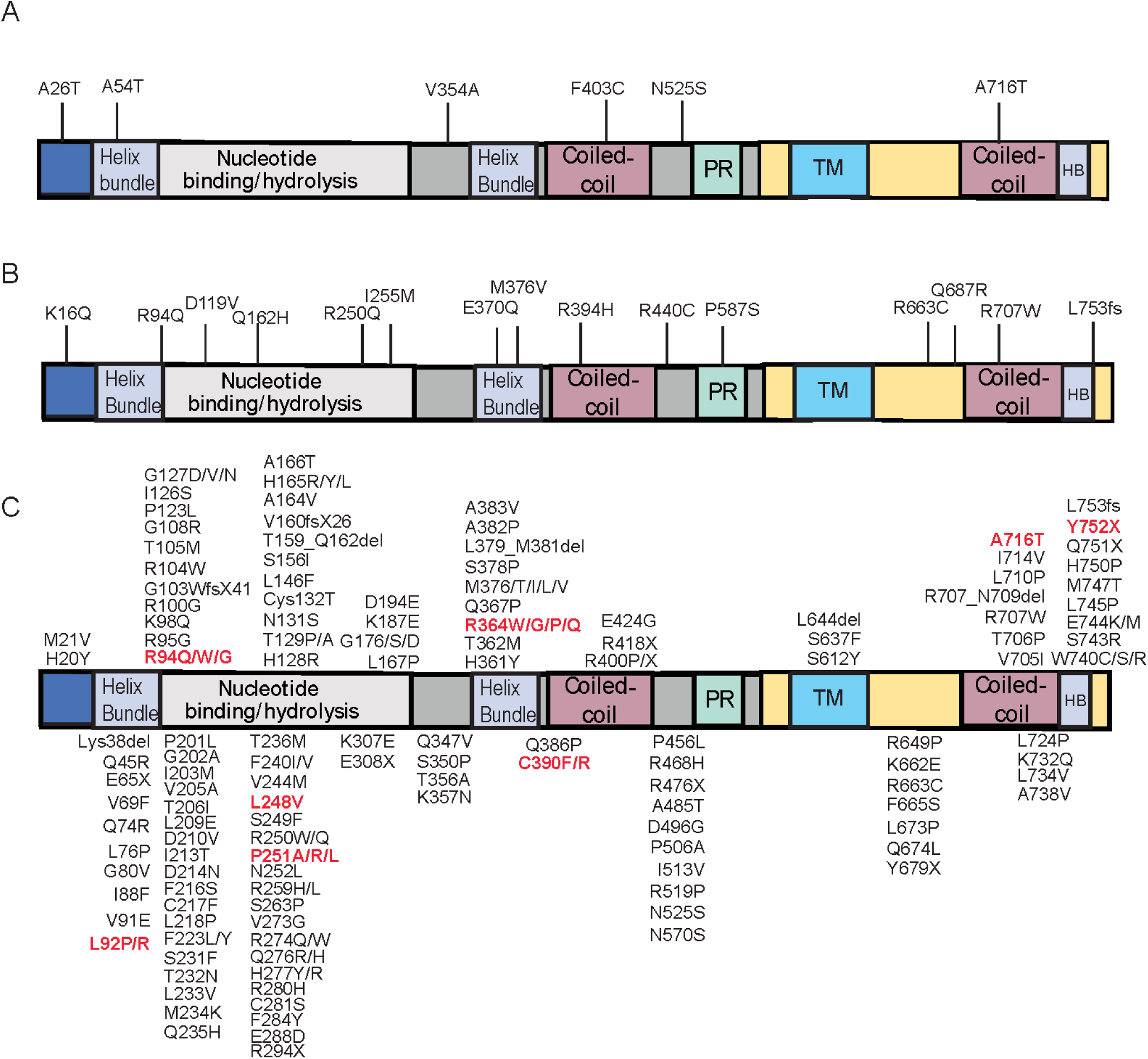
A) Schematic of MFN2 protein domains and six mutations initially identified in the University of Utah ALS cohort. B) Fifteen additional MFN2 mutations identified in ALSdb. C) Reported mutations in MFN2 that cause CMT2A. Variants shown in red have been associated with predominantly motor phenotypes in CMT2A. HB=helix bundle, PR=proline rich domain, TM=transmembrane.

## METHODS

### VAAST and Phevor utilization to identify candidate genes

To identify genes that were more burdened with rare, deleterious variants in SALS cases compared to controls, we utilized VAAST (Variant Annotation, Analysis, and Search Tool) and Phevor (Phenotype Driven Variant Ontological Re-ranking) software^15,16^. VAAST determines gene burden by assessing both variant rarity and the mutation severity. VAAST is an aggregative approach to determine the overall effect of mutations on gene function. VAAST output is then processed by Phevor, which re-ranks candidate loci based upon ontological data (such as the Disease Ontology and Gene Ontology databases). Our pipeline and the ALS cohorts we utilized have been described previously^4,17^. In total, we analyzed sequence data for a total of 140 ALS patients.

### Determining allele frequency for identified mutations

To determine the frequency of the mutations identified in our ALS cohort, we converted the GRCh37 (hg19) coordinates to GRCh38 (hg38) coordinates using LiftOver (UCSC Genome Browser). GnomAD 3.1 contains 76,156 whole-genome sequenced individuals from diverse backgrounds and disease cohorts. To best match the genetic background of the cases in this study, we used the gnomAD 3.1 VCF to find the minor allele frequencies for each of the mutations identified in MFN2 in the Non-Finnish European control/biobank cohort (n = 3,427). We repeated this process in the control dataset included in gnomAD 2.1, which also includes exome sequencing data. For variants in gnomAD 2.1 we did not use liftOver to convert the coordinates, as the variants are listed with GRCh37 coordinates.

### *Mfn2* KO MEF cell culture and mutant MFN2 lentiviral production

*Mfn2* KO mouse embryonic fibroblasts (MEFs)^6^ were compared with wildtype control MEFs. MEFs were cultured in DMEM high glucose with glutamate and 10% fetal bovine serum added. The Tet-on system was used to induce gene expression using pCLX-pTF-R1-DEST-R2-EBR65 (addgene #45952). A doxycycline concentration of 100ng/mL was used and was refreshed every 48 hours. Cells were on doxycycline for 72 hours before imaging or staining. Wildtype coding sequence of *MFN2* was amplified from MFN2-YFP (a gift from Richard Youle Addgene#28010). Mutations found in ALS patients were introduced using site-directed mutagenesis and cloned into pDONOR221. Next, LR reactions between pDONOR221 and pCLX-pTF-R1-DEST-R2-EBR65 were performed. HEK293T cells were transfected using calcium phosphate with psPAX2 (gift from Didier Trono, Addgene No.12260), pCAG-Eco (gift from Arthur Nienhuis and Patrick Salmon, Addgene No. 35617), and cloned pCLX-DEST-MFN2 vectors. Supernatant was harvested 72 hours later for infection of KO MEFs.

### Western blotting of infected *Mfn2* KO MEFs

To confirm mutant protein expression, MEFs expressing mutant MFN2 were exposed to dox for 72 hours. After 72 hours, cells were lysed using RIPA buffer plus protease inhibitor tablets. 40ug of protein were loaded in 10% Bis-Tris gels and run with MOPS buffer. Gels were transferred onto PVDF membranes (preactivated with methanol) for 40 minutes on ice at 200mA. Blots were blocked for one hour in 5% milk/TBST. Blots were then incubated overnight at 4 degrees with antibodies: rabbit anti gapdh (1:2000, ProteinTech 10494-1-AP), mouse anti MFN2 (1:200, Novus# H00009927-M01). Secondary antibodies were goat anti-rabbit or mouse conjugated to HRP (Thermo). Developing solution was clarity western ECL (BioRad).

### Mitochondrial morphology and membrane potential in mutant-MFN2 expressing KO MEFs

Cells were seeded onto sterile glass slides and were exposed to dox for 72 hours. Cells were fixed with 4% paraformaldehyde for 15 minutes, washed with PBS, and permeabilized with PBS/0.002%TritonX. Cells were washed, blocked for one hour with 10% goat serum, then incubated with 1:200 primary antibody (mouse anti ATPB abcam #AB14730) for 1.5 hours. Cells were washed, then incubated with 1:200 goat anti mouse-Alexa 488 (Thermo) for 45 minutes, washed, then mounted with Prolong Diamond. Leica SP8 confocal was used with 0.2 um step size for 3D rendering. To determine mutant rescue efficiency, mitochondrial morphology for each cell image was binned into one of three categories: fused, intermediate, or fragmented. We were blinded to genotype. Four replicates of 100 cells each were analyzed for the original six mutations (x1-x6) in MFN2 identified in our University of Utah ALS cohort. Two replicates of 100 cells each were analyzed for each ALSdb mutation. To determine significance, a Fisher’s Exact test was used with categories classified as Fused vs non-fused (intermediate and fragmented were combined since it has to be a 2×2 table). After blinded analysis, images were run through Mitochondria Analyzer (MA) software (v1) to analyze mitochondrial metrics in ImageJ^18^.

For membrane potential analysis, cells were seeded in ibidi 8uwell plates (#80826) and then exposed to dox for 72 hours. Cells were incubated with 150 nM TMRE (Biotium, 70005) for 15 minutes. Cells were washed with PBS and then incubated with FluoroBrite DMEM (Thermo) for live cell imaging on Leica SP8. TMRE intensity was quantified in Fiji.

### Genotyping, movement assay, and protein analysis of *mfn2* KO zebrafish

Zebrafish genotyping was performed using nested PCR to introduce PsiI-v2 cut sites, with enzymatic cutting being dependent on the L285X mutation^19^. DNA was amplified using Qiagen PCR Taq (#201205). At 48 hpf, fish were touched on their tail. Movement was quantified based on whether the fish swam away, jerked in place, or did not move at all. We were blinded to genotype and 243 fish were analyzed. Mutant fish were iced and then incubated in 4% paraformaldehyde overnight at 4 degrees. Fish were dehydrated in 100% methanol and stored at -20 until later use. Zebrafish brains were dissected into four sections: forebrain, hindbrain, cerebellum, and optic tectum. Proteins were extracted using Qiagen Qproteome FFPE tissue kit and DNA extracted with QIAamp DNA FFPE tissue kit. Rabbit anti β-Tubulin (1:5000 CST #2146S), rabbit anti-GAPDH (ProteinTech), and rabbit anti-TDP-43 (1:200, ProteinTech #10782-2-AP) were the primary antibodies used with subsequent use of HRP-conjugated secondary antibodies.

## RESULTS

Analysis of whole-genome sequencing data from 140 ALS patients revealed two top candidate loci, *TP73* and *MFN2*^17^. Six rare (minor allele frequency < 0.001), nonsynonymous, deleterious mutations in *MFN2* were identified (Fig1A). We also screened the publicly available database ALSdb, composed of ∼2,800 ALS patients, in which we identified 15 additional rare, deleterious mutations in *MFN2* (Fig1B).

We performed rescue experiments to determine whether these mutations in MFN2 have a deleterious effect on protein function. Loss of Mfn2 causes mitochondria to become fragmented and to appear as spherical puncta (Fig 2). We cloned the *MFN2* coding sequence and all 21 mutations into lentiviral vectors for infection of KO MEFs in a total of 21 mutant-expressing lines (Methods). As controls, two distinct point mutations that encode S523S and V705I were inserted into the wildtype coding sequence of MFN2. S523S is the most common synonymous mutation in *MFN2* in the gnomAD database and V705I is the most common nonsynonymous mutation in *MFN2* in gnomAD(v2). These were expressed in KO MEFs to show that creating a SNP in the *MFN2* coding sequence did not inhibit rescue and that non-rescue would be due to the specific mutations identified in ALS patients. As a control, we cloned the coding sequence for eGFP into the lentiviral vector for infection.

**Fig 2.**
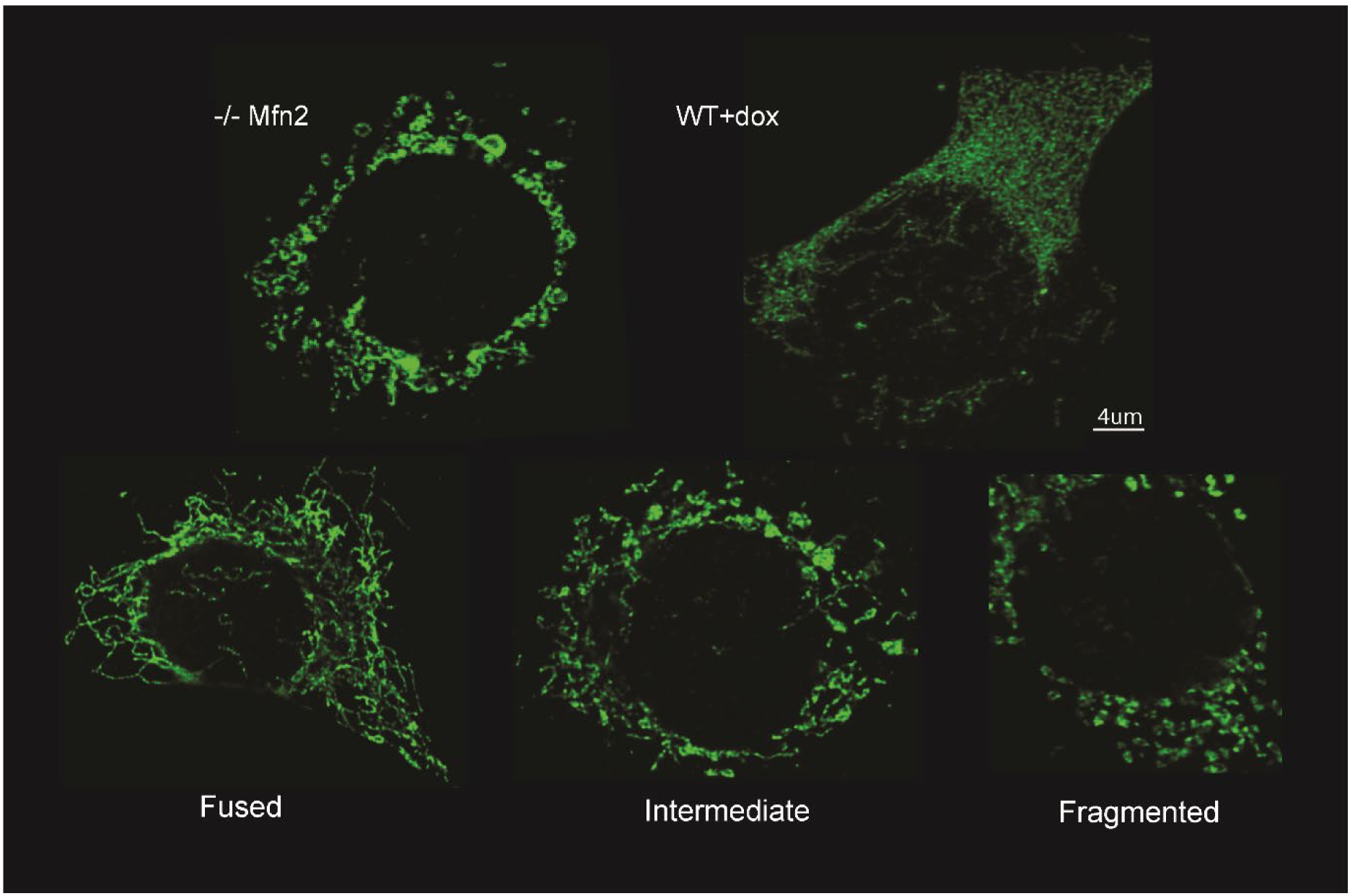
Confocal images of *Mfn2* KO MEFs, wildtype (WT) MEFs exposed to dox, and the three categories that mutant-expressing KO MEFs were binned into during morphological analysis. Mitochondria are stained green.

We utilized the Tet-on system, where doxycycline exposure is needed for gene expression. After induction with doxycycline, staining for mitochondria, and confocal imaging, morphology was analyzed with blinding to genotype. Mitochondrial morphology was binned into one of three distinct categories: fused, with long, branched networks; intermediate, with less complex networks; or fragmented, with most mitochondria displaying spherical puncta (Fig2). Wildtype MEFs, MEFs + dox, KO MEFs expressing MFN2WT (KO + MFN2WT), KOs expressing MFN2S523S (KO + MFN2S523S), KOs expressing MFN2V705I (KO + MFN2V705I), KO MEFs, and KO MEFs expressing eGFP (KO + GFP) were scored. These controls and the original six mutations (denoted x1,x2,x3,x4,x5,x6) identified in the University of Utah ALS cohort were scored from 4 replicates of 100 cells each for a total of 400 cells (Fig3,5). The 15 ALSdb mutants were scored from 2 replicates of 100 cells each for a total of 200 cells (Fig4,5).

**Fig 3.**
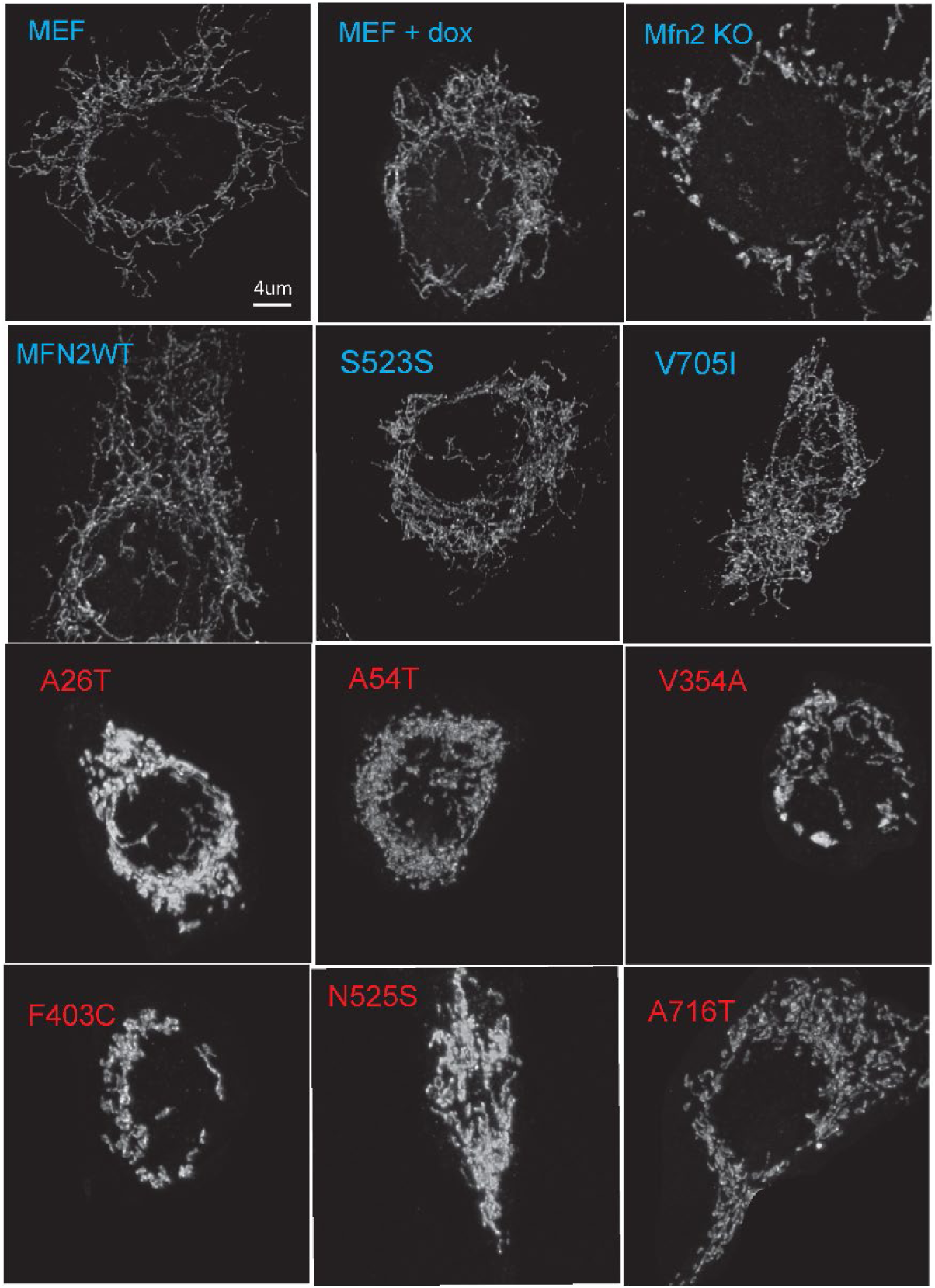
Sample confocal images of mitochondria in controls and mutants expressing KO MEFs. Control lines are in blue and original six mutations in ALS patients are in red.

**Fig 4.**
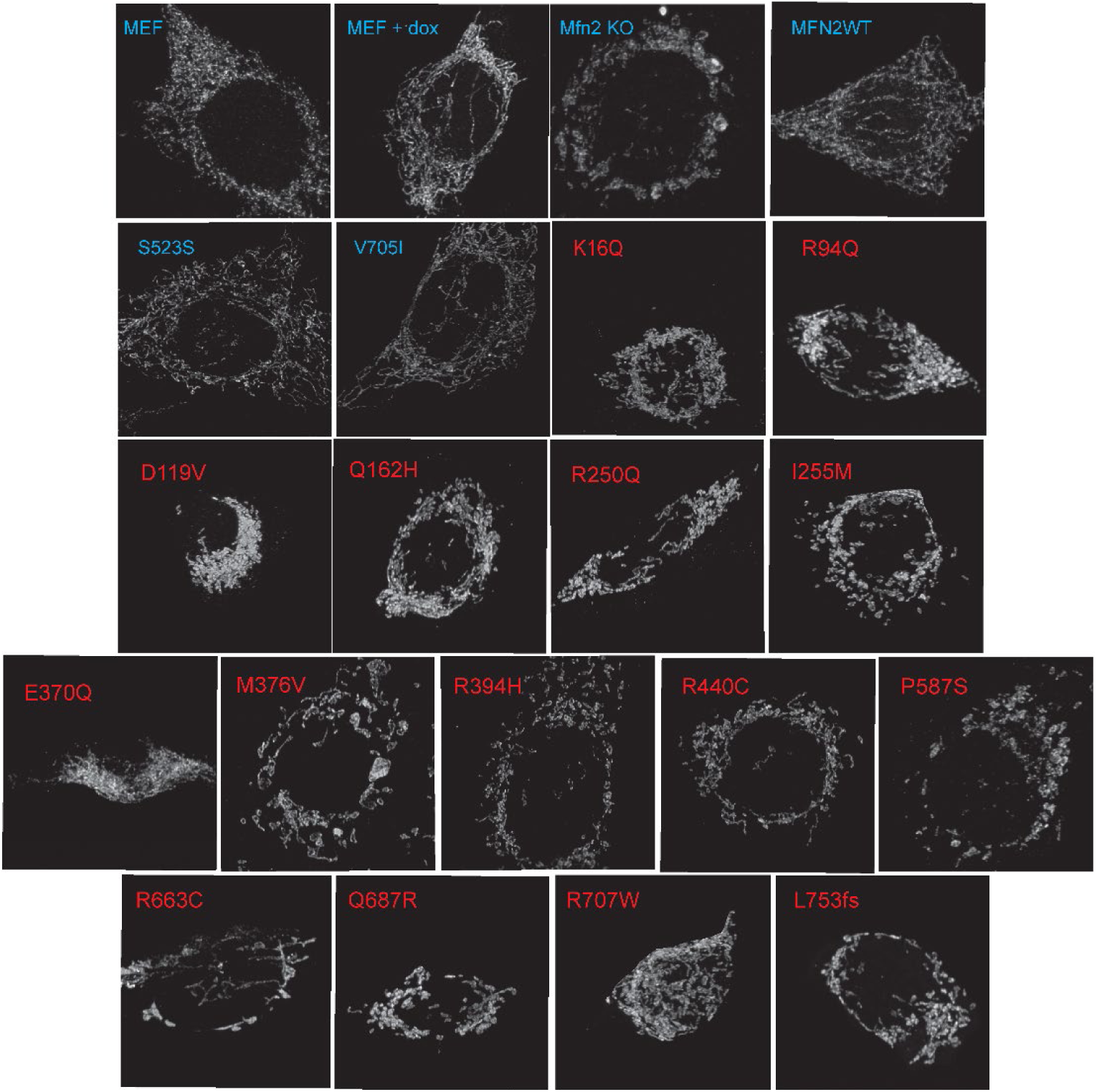
Sample confocal images of mitochondria in controls and mutants expressing KO MEFs. Control lines are in blue and the 15 ALSdb mutants are in red.

**Fig 5.**
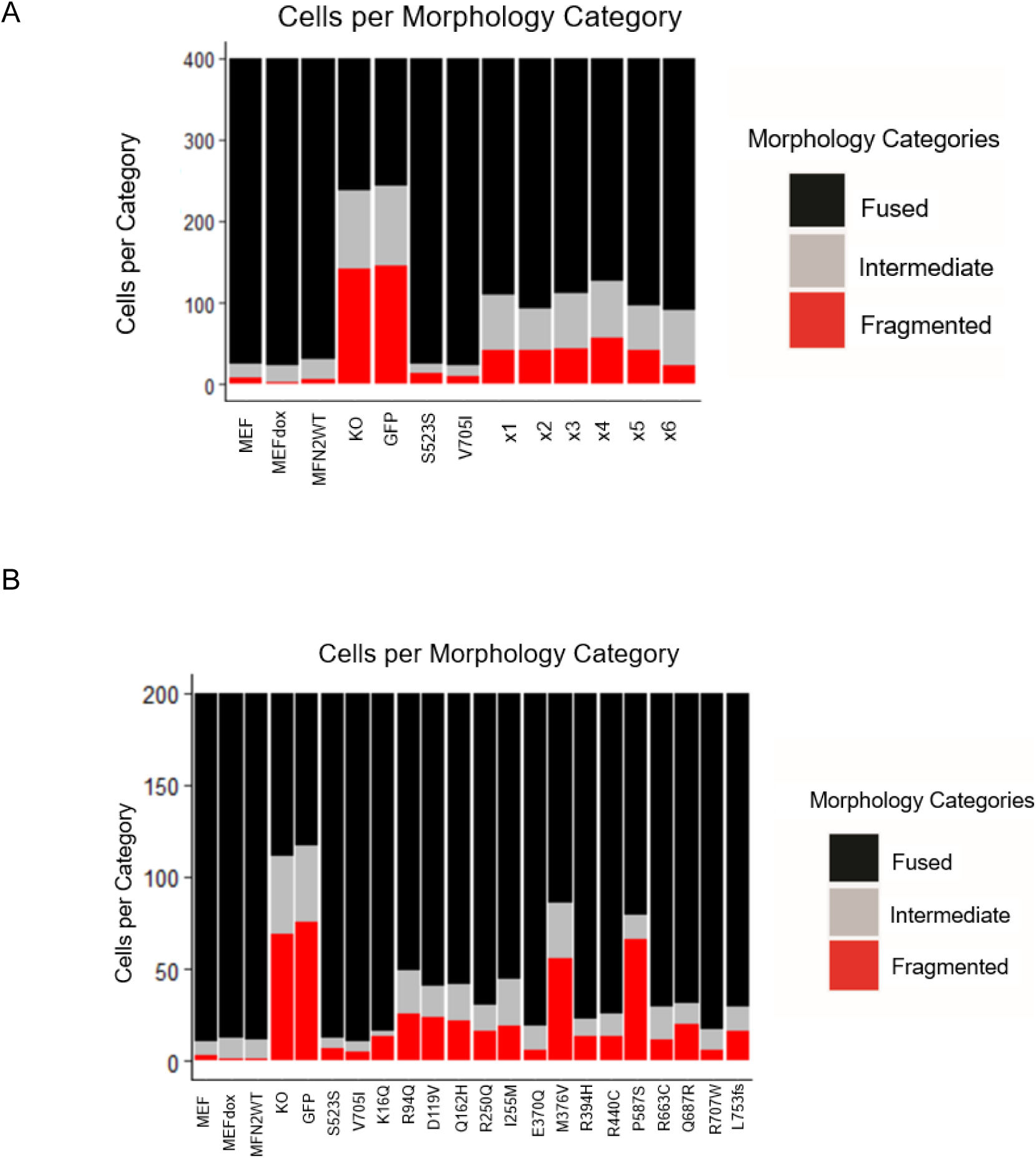
A) Stacked bar graph showing how many cells were classified into each mitochondrial morphology category for x1-x6, which represent the original six mutations identified in MFN2. B) Stacked bar graph showing number of cells per morphological category for KOs expressing the fifteen ALSdb mutations. S523S and V705I are the gnomAD control variants.

Wildtype MEFs, MEFs + dox, KOs + MFN2WT, KOs + MFN2S523S, and KOs + MFN2V705I showed few cells with intermediate or fragmented mitochondria. The vast majority of these cells had elongated, fused mitochondria, as expected (Fig.4). When Mfn2 KO MEFs and KOs + GFP cells were scored, the number of cells showing fragmented and intermediate mitochondrial morphologies increased drastically. This was expected, as this was the morphological phenotype observed when the *Mfn2* KO MEFs were first generated^6^. The initial six mutations (labeled x1-x6) identified in MFN2 in ALS patients all showed a significant increase in the number cells with fragmented and intermediate mitochondrial morphologies (Fisher’s exact test, p<0.005, Fig5A). Mutant x4 showed the largest increase in the number of cells with fragmented mitochondria. Mutants x1, x3, x4, and x6 showed similar increases in the number of cells with intermediate mitochondrial morphology. ALSdb mutants R94Q, Q162H, I255M, and M376V showed the largest increase in the number of cells with intermediate mitochondrial morphology(Fig5B). R94Q, D119V, Q162H, I255M, Q687R, and L753fs showed similar increases in the number of cells with fragmented mitochondria. The largest increases were seen in mutants M376V and P587S. The most severe mutants in terms of rescue efficiency were x1, x2, x3, x4, x5, R94Q, D119V, Q162H, I255M, M376V, P587S, Q687R, and L753fs (frameshift).

To test our hypothesis that the mutations cause insufficient rescue, we next used Mitochondria Analyzer (MA)^18^, which quantitatively analyzes 2D/3D images of mitochondria. When mitochondrial count for KO MEFs expressing wildtype MFN2 was compared to mitochondrial count for MEF, MEF + dox, Kos +MFN2S523S, and KOs + MFN2V705I, there was no significant difference (Fig 6-7). This shows that rescue with wildtype MFN2 restores mitochondrial count to that of MEFs and that inserting common SNPs from gnomAD did not affect rescue. As expected, when comparing KO + wildtype MFN2 to *Mfn2* KO MEFs, mitochondrial count was significantly decreased. Count was significantly decreased when comparing KO + MFN2WT to KOs expressing mutations x1-x6, with x1, x2, and x5 showing the most significant decrease. When comparing KO + MFN2WT counts to the 15 mutations identified from ALSdb, there was a statistically significant decrease for all mutants (Wilcoxon-Mann-Whitney test, multiple comparisons corrected using the Holm method, p<0.0001). The smallest decreases were seen for mutants Q162H, E370Q, P587S, and R707W. The largest decreases in mitochondrial count were seen for mutants R94Q, R250Q, R394H, D119V, M376V, R440C, R663C, Q687R, and L753fs. Similar trends were seen when branch number was quantified (not shown).

**Fig6.**
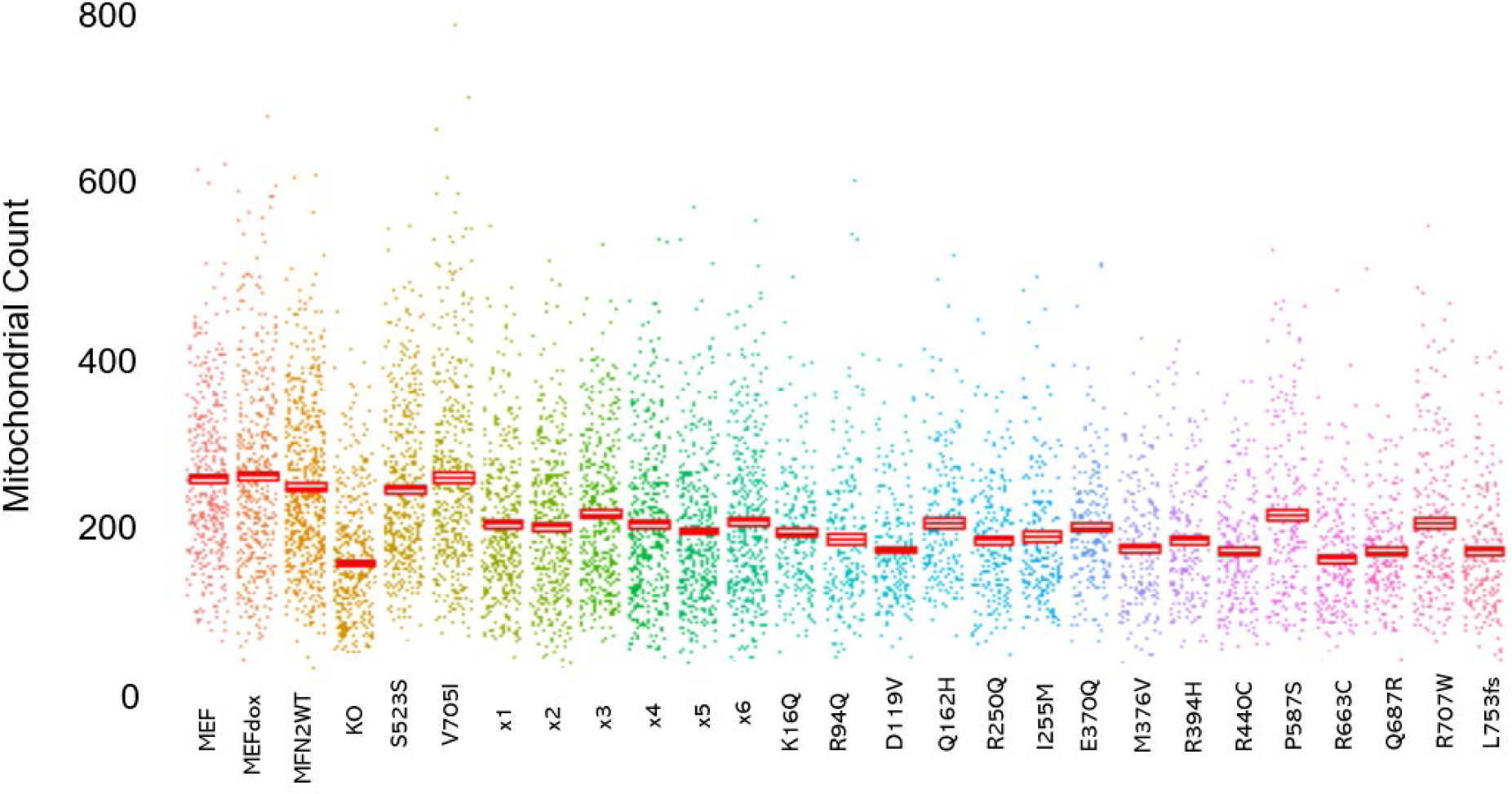
Quantification showing mean +/- standard error (SE) for mitochondrial count quantified by MA software^19^. X axis shows the genotype and y axis shows mitochondrial count. S523S and V705I are the control gnomAD variant mutations, x1-x6 are the original six mutations identified in MFN2, followed by the fifteen mutations from ALSdb.

**Fig7.**
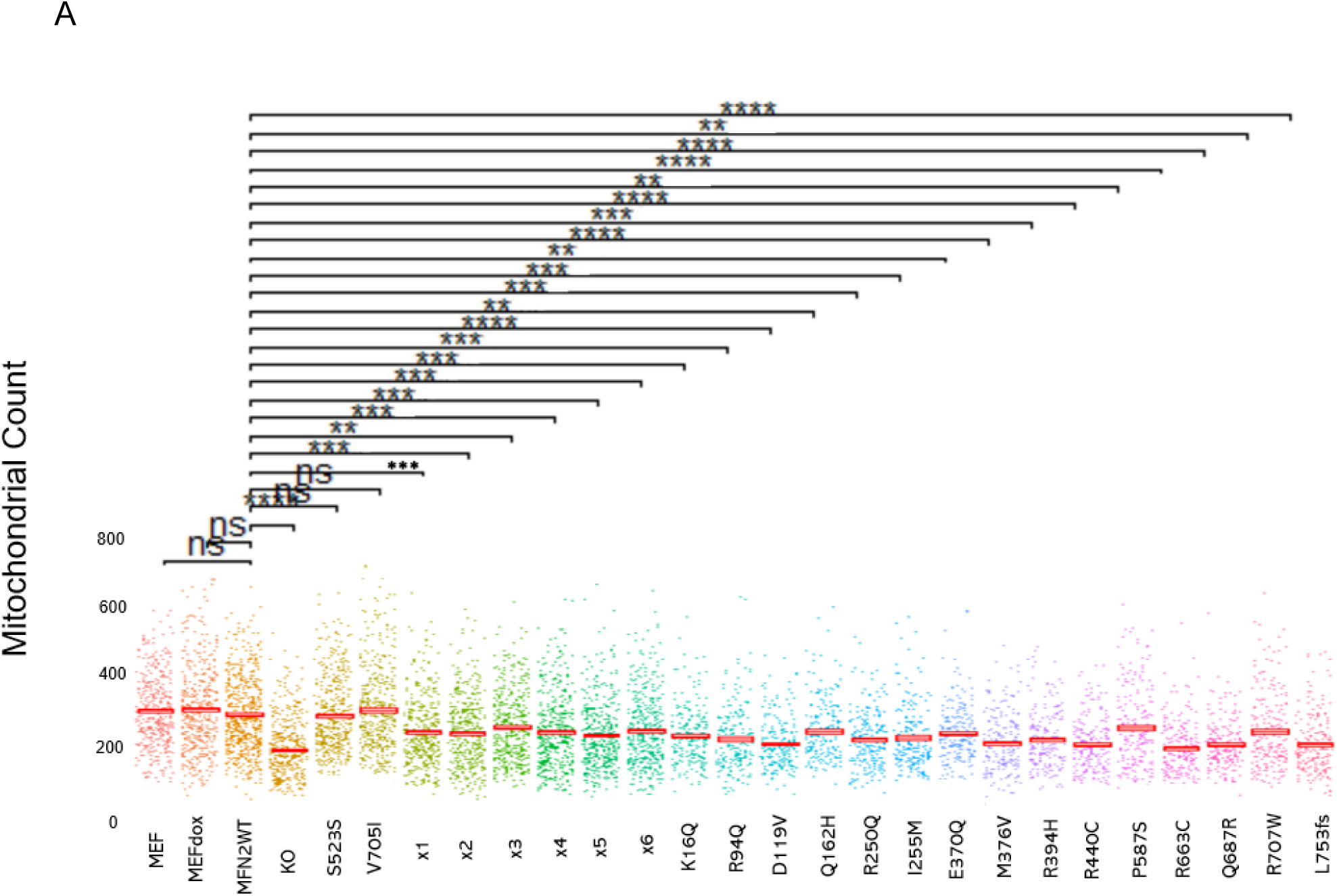
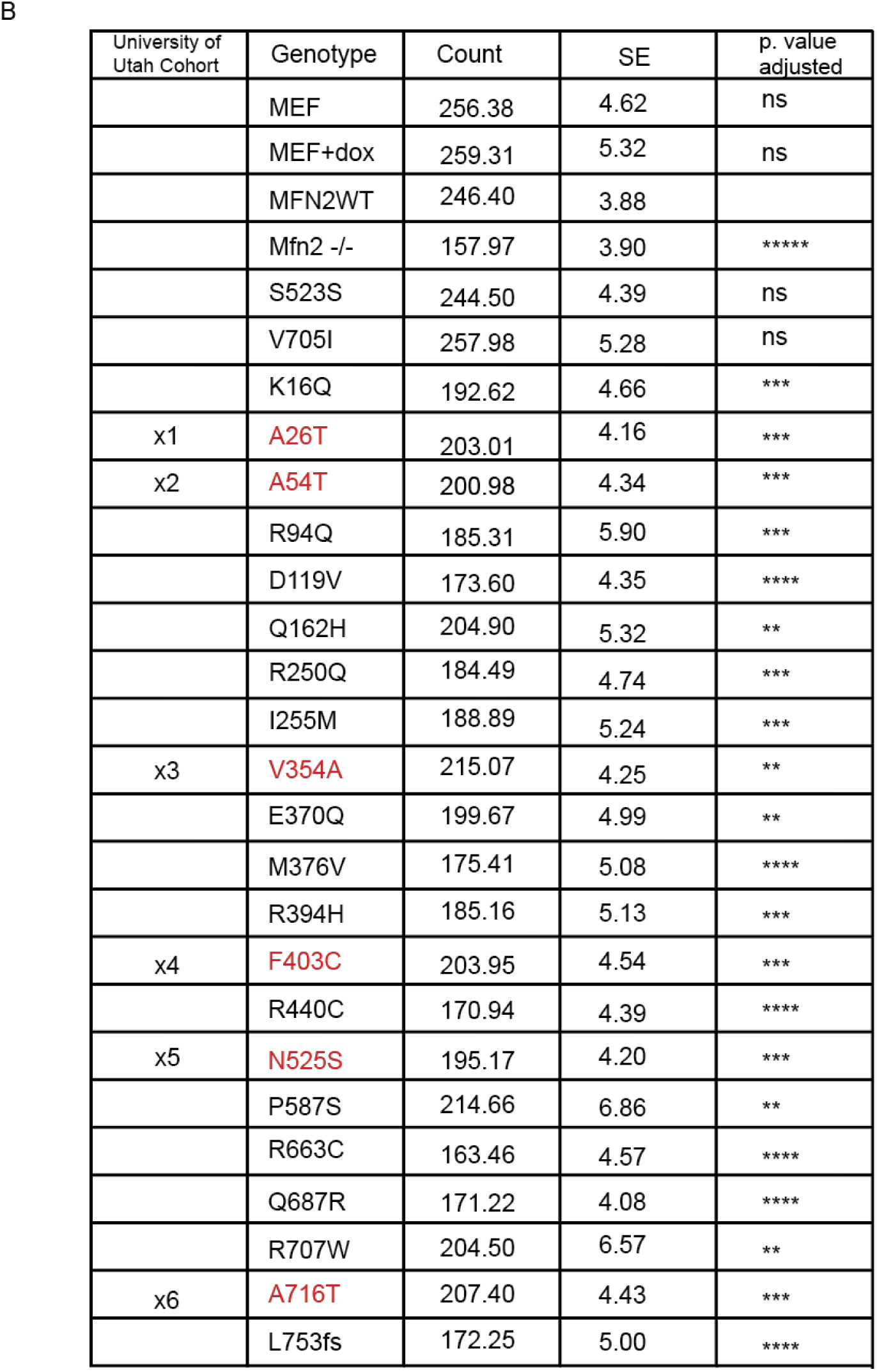
A) Statistical comparison between KO + MFN2WT compared to all genotypes. B) Mean, SE, and corrected p values for mitochondrial count from MA analysis. Wilcoxon-Mann-Whitney, Holm correction for multiple comparisons, **p<0.0001, ***p<1^-10^, ***p<1^-18^

These results show that the mutations we identified in MFN2 in ALS patients render the wildtype protein dysfunctional. Expression of S523S and V705I showed no difference in any parameters when compared to KO + MFN2WT, demonstrating that the patient-derived mutations have an *in vitro* effect and cause insufficient rescue.

Next, we measured membrane potential of KOs expressing x1-x6 and gnomAD control variants. All mutants showed significantly decreased membrane potential compared to MFN2WT, S523S, and V705I expression, except for x6 (p<0.00001). This defect in membrane potential strengthens our argument that these mutations have a deleterious effect on mitochondrial function, but more membrane potential experiments are needed.

To study loss of *mfn2 in vivo* and to determine whether these KO animals could be used as a novel ALS model, we studied *mfn2* KO zebrafish^19^. Crosses of *mfn2* +/-heterozygous adults produced embryos with morphological defects such as curved tails, like those seen in mutant TDP-43 zebrafish (Fig8A)^20^. Genotyping revealed that these morphologically deformed embryos were heterozygous or homozygous for loss of *mfn2*. We also investigated if young fish had movement defects. At 48 hpf, fish were touched on their tail and their movement was quantified (Fig8B). We saw a significant increase in the proportion of heterozygous and homozygous fish that jerked in place/did not swim away compared to wildtype. Heterozygous fish also showed an increase in no movement upon touch compared to wildtype. We report that mutant fish have movement and morphological abnormalities as early as 48 hpf.

**Fig.8.**
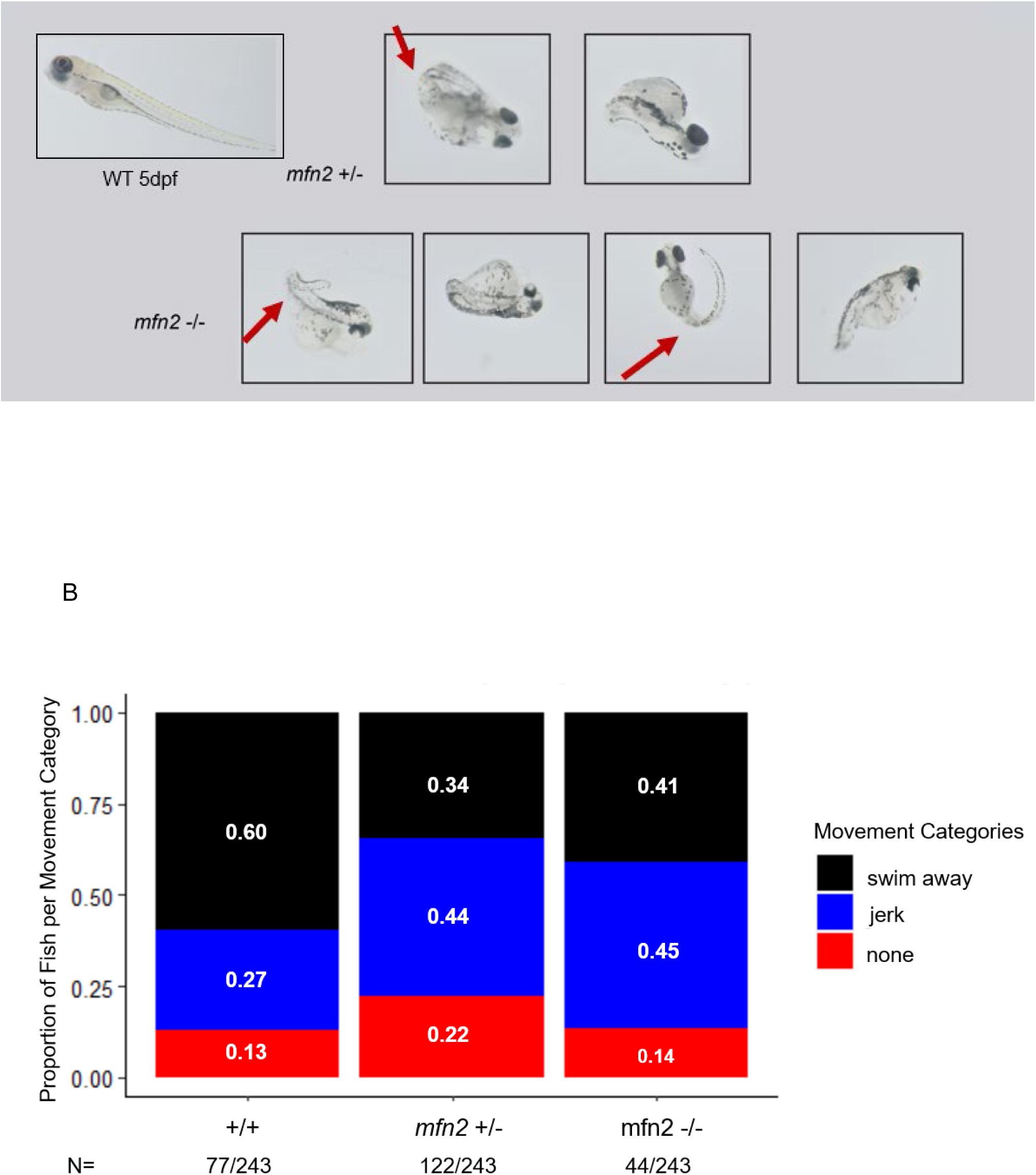
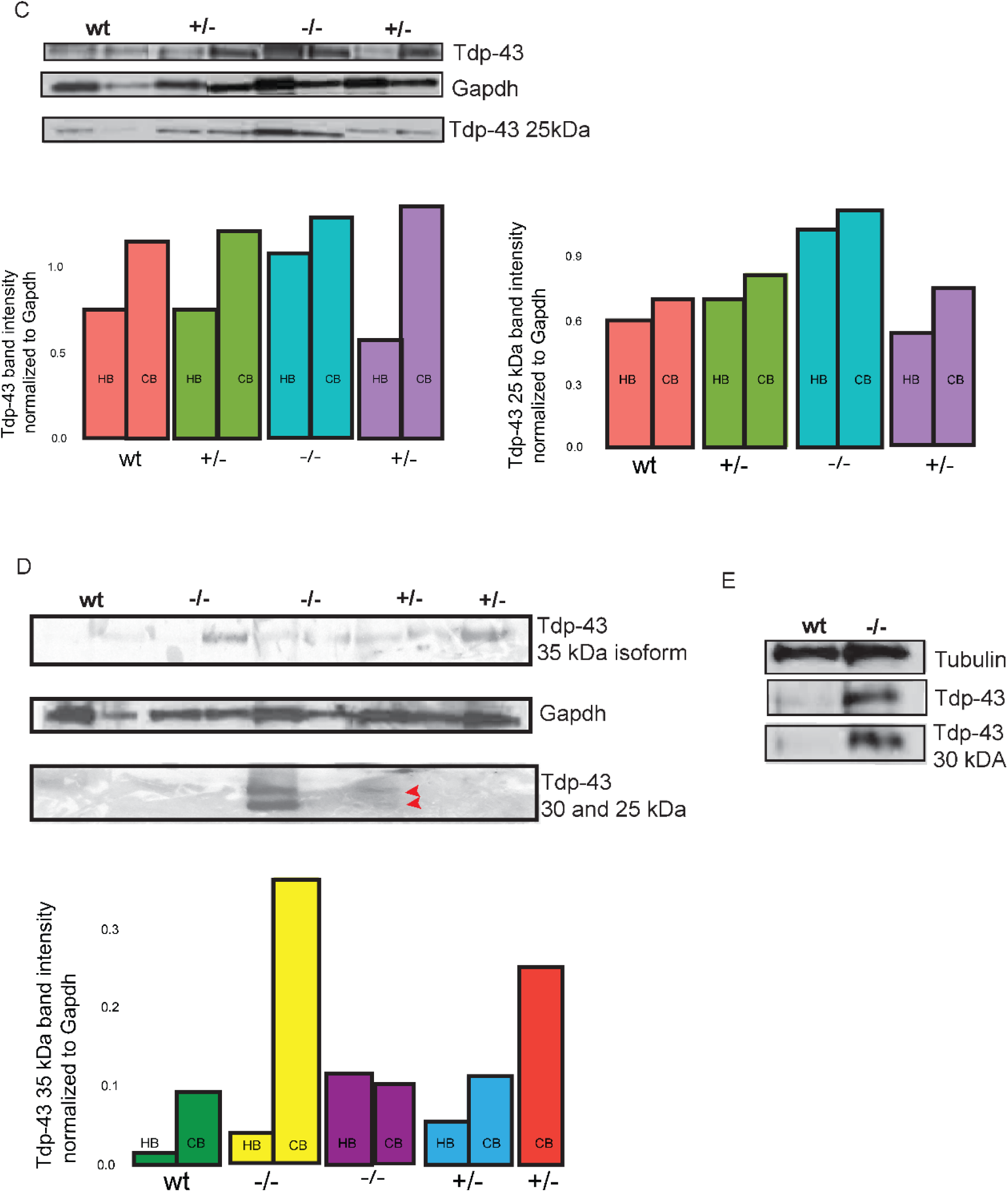
A) Heterozygous and homozygous mutant zebrafish show curly tails as early as 5 dpf (red arrows), as in other zebrafish ALS models^20^. B) Blinded to genotype, 48 hpf zebrafish were binned into three different categories based on movement after tail touching. White numbers in the stacked bar graph are the proportion of fish per genotype represented in that category. C-E) Western blots of brain lysates of aged wildtype, +/-, and mfn2 -/-fish. C) Mutant fish show increases in Tdp-43 and its smaller isoform at 25 kDa. Quantification of band intensity normalized to Gapdh are below the blot. D) Mutant fish show increases in Tdp-43 isoforms at 35, 30, and 25 kDa not seen in wildtype. Red arrows show faint aggregation at 30 and 25kDa in heterozygous brain lysate. Quantification is below the blot. E) Western blot illustrating the significant increase in Tdp-43 and smaller isoform at 30 kDa in mutant fish not seen in wildtype. HB= hindbrain, CB=cerebellum, wt = wildtype, +/-= heterozygous mutant, -/-= homozygous mutant.

Aged, mutant zebrafish were fixed and their brains dissected to determine if TDP-43 aggregation was occurring, which is a hallmark of ALS pathogenesis^20^. In brain lysates for the hindbrain and cerebellum of heterozygous and homozygous *mfn2* mutant fish, we saw increases in TDP-43 (Fig8B,C). We also saw increases in isoforms of TDP-43, at 35, 30, and 25 kDA. Isoforms of these sizes have been reported in ALS pathology^20,21.^ These increases were not seen in wildtype brain lysates. In summation, our *in vitro* and *in vivo* results show that ALS-patient mutations in MFN2 cause protein dysfunction, and loss of *mfn2 in vivo* is a novel model of ALS.

## DISCUSSION

There is increasing evidence that dysfunctional mitochondria may be an important component of ALS pathology. Mitochondrial defects have been seen in many *in vitro* studies of ALS and in multiple ALS rodent models^22-35^. In addition, inhibition of mitochondrial fission (through Drp1) can slow the progression of ALS^32^. CHCHD10 is an ALS-causing, mitochondrial protein, and marf (drosophila ortholog of MFN2) has been suggested to be a downstream effector of CHCHD10-mediated damage^36^. In SOD1 mice, swollen mitochondria located in vacuoles with atypical cristae organization were observed early in disease pre-symptom onset^37^. Multiple studies have shown that TDP-43’s neurotoxicity is due to its negative impact on mitochondrial function^27-31^. In normal conditions, TDP-43 is predominantly found in the nucleus, but TDP-43 is also localized with mitochondria^31^. Mice expressing mutant TDP-43 demonstrated disorganized mitochondria, and expression of TDP-43 in primary neurons caused mitochondrial fragmentation, slowed mitochondrial transport, and decreased membrane potential ^27,28^. Importantly, Mfn2 overexpression prevented this fragmentation, restored mitochondrial length and mitochondrial transport rates, and reversed membrane depolarization induced by mutant TDP-43. These mice also had abnormal accumulations in their brainstem and spinal motor neurons that consisted of grouped mitochondria. Additionally, expression of Mitofusin 1 was decreased in these mutant mice while Fis1 and phosphorylated Dlp1 (pro-fission factors) were both upregulated.

In ALS patient spinal cord or frontotemporal dementia (FTD) cortical neuron samples, it was determined that mislocalized, cytoplasmic TDP-43 was mostly associating with mitochondria, specifically the inner mitochondrial membrane (IMM)^30^. At much lower levels, this was also seen in control samples, showing that part of TDP-43’s normal function is to interact with mitochondria. Mutations in TDP-43 that cause ALS (G298S or A3382T) caused increased TDP-43 association with mitochondria with increased import into the IMM, yet there were no changes in overall total protein amounts of TDP-43. Giving perhaps an insight into TDP-43’s normal function in mitochondria, TDP-43 was shown to bind mitochondrial transcribed mRNAs^30^. Additionally, knocking down TDP-43 in HEK293T cells led to a significant decrease in MFN2 protein^29^. It was subsequently determined from human cortical brain tissue that TDP-43 is a binding partner of MFN2. Importantly, blocking TDP-43 from associating with mitochondria prevented its neuronal toxicity and significantly improved neuromuscular junction loss/motor-defects in TDP-43 mutant mice^30,31^. Highlighting MFN2’s importance, overexpression of marf in *Drosophila* alleviated the disease phenotype induced by expression of TDP-43^35^.

A short tandem repeat expansion of a hexanucleotide tandem repeat within *C9orf72* is the most common known cause of ALS, and recently it was shown that the encoded C9orf72 protein maintains the respiratory chain of mitochondria^23^. C9orf72 was found to reside in the IMM, and C9orf72 KO MEFs had a significant decrease in oxidative phosphorylation. Upon stress exposure, the loss of C9orf72 caused decreased ATP production compared to WT (specifically through inhibited complex I activity). This was verified by a significant decrease in tetramethylrhodamine, methyl ester (TMRM) intensity, showing that loss of C9orf72 leads to aberrant mitochondrial function. Dysfunctional transport of mitochondria along axons was observed in motor neurons differentiated from C9orf72 expansion patient samples^24^. There was a decrease in motile mitochondria, and mitochondrial respiration was irregular. This seemed to be due to a decline in mitochondrial transcripts encoded by mitochondrial, but not nuclear, genes. Perhaps C9orf72 is affecting mitochondrial mRNAs in a manner similar to TDP-43. These studies all highlight the importance of mitochondria, and MFN2, in neuronal health and ALS causation.

MFN2 protects against glutamate excitotoxicity, a process in which the overexcitation of neurons by glutamate leads to mitochondrial fragmentation, decreased MFN2 protein levels, and motor neuron demise^38^. Riluzole, the only broadly available treatment for ALS, is thought to help prevent glutamate excitotoxicity. MFN2 overexpression in primary motor neurons prevents glutamate-induced neuronal death^38^. Perhaps the neurons of patients with mutant MFN2 are more susceptible to glutamate-induced death due to abnormal MFN2 function, which eventually leads to ALS. MFN2 has also been shown to protect neurons against other forms of stressors^39,40^. Both constitutively active and wildtype Mfn2 overexpression in primary cerebellar granule neurons protected against neuronal death induced by DNA damage or reactive oxygen species through mitigation of cytochrome c release. Even when no cell death induction reagents were used, knock-down of Mfn2 caused one-third of neurons to die. Mfn2 has also been implicated in autophagy, a pathway heavily implicated in ALS^41,42^.

We propose that mutations in MFN2 influence ALS pathology and that MFN2 is a novel modifier of disease phenotype. To our knowledge, this is the first report of mutations in *MFN2* associated with ALS. Multiple lines of evidence support a role of MFN2 in ALS pathology. First, we show that twenty-one distinct mutations in MFN2 identified in ALS patients render the protein defective in rescuing morphological defects in *Mfn2* KO MEFs. Second, overexpression of MFN2 has been shown to protect against TDP-43-induced mitochondrial dysfunction in mouse neurons^27^. Third, MFN2 also protects against glutamate excitotoxicity, a pathway known to be involved in ALS pathogenicity^38^. MFN2 has also been shown to be necessary for cerebellar development in mice, and any deleterious mutations in MFN2 could leave neurons more vulnerable to stress/insults^8^.

Fourth, MFN2 causes the sensory and motor neuropathy disease CMT2A. Multiple animal models have shown that loss of *Mfn2* function leads to abnormal motor neuron morphology and movement defects^7,8,43,44^.

There have been multiple reports of CMT2A patients who present with predominantly motor phenotypes^9-14^. Some of the mutations in *MFN2* we discovered in ALS patients have been reported in CMT2A pathogenesis, and R94Q is one of the most common causes of CMT2A^10^. However, most of the mutations we report in MFN2 in ALS patients have not been reported in CMT2A. These mutations may be more severe and/or have a different mode of action that leads to ALS instead of CMT2A. Factors such as genetic background could influence disease progression towards either ALS or CMT2A as well. We propose that CMT2 and ALS lie on a disease continuum, as ALS does with FTD. Known ALS genes *CHCHD10* and *FIG4* cause forms of CMT, as we propose for MFN2^45,46^.

Lastly, we utilized previously characterized *mfn2* KO zebrafish to determine whether these fish developed ALS-like symptoms^19^. Mutant embryos would develop a curly tail phenotype, mutant fish display movement abnormalities at 48 hpf, and cerebellum/hindbrain lysates displayed increased levels of TDP-43 (at its normal size and isoforms at lower molecular weights) in heterozygous and homozygous fish. Interestingly, we observed that some, but not all, adult heterozygous fish had motor and size defects similar to homozygous mutants. This suggests that haploinsufficiency of *mfn2* could potentially lead to motor neuron disease *in vivo*. MFN2 has been shown to be a druggable target in neurodegeneration^43^. Our preliminary results raise exciting possibilities for therapeutic drug screening, augment knowledge of mitochondrial function in ALS pathology, and highlight that *mfn2* KO zebrafish can be used as a novel ALS/motor neuron disease model.

